# Ultrapotent and Broad Neutralization of SARS-CoV-2 Variants by Modular, Tetravalent, Bi-paratopic Antibodies

**DOI:** 10.1101/2021.11.02.466984

**Authors:** Shane Miersch, Reza Saberianfar, Chao Chen, Nitin Sharma, Gaya K. Amarasinghe, Francesca Caccuri, Alberto Zani, Arnaldo Caruso, Giuseppe Novelli, Sachdev S Sidhu

## Abstract

Neutralizing antibodies (nAbs) that target the SARS-CoV-2 spike protein are approved for treatment of COVID-19. However, with the emergence of variants of concern, there is a need for new treatment options. We report a novel format that enables modular assembly of bi-paratopic, tetravalent nAbs with antigen-binding sites from two distinct nAbs. The tetravalent nAb was purified in high yield, and it exhibited biophysical characteristics that were comparable to those of approved therapeutic antibodies. The tetravalent nAb bound to the spike protein trimer at least 100-fold more tightly than bivalent IgGs (apparent K_D_ < 1 pM), and it exhibited extremely high potencies against a broad array of pseudoviruses, chimeric viruses, and authentic virus variants. Together, these results establish the tetravalent diabody-Fc-Fab as a robust, modular platform for rapid production of drug-grade nAbs with potencies and breadth of coverage that greatly exceed those of conventional bivalent IgGs.

## Introduction

SARS-CoV-2, the causative agent of the ongoing COVID-19 pandemic, relies on its surface spike glycoprotein (S-protein) to mediate interaction with and entry into host cells (Hoffmann et al., 2020). The surface of the virus particle is studded with 25-100 copies of S-protein homotrimers, and each S-protein consists of two subunits (Klein et al., 2020; Walls et al., 2020). The N-terminal subunit (S1) mediates host cell recognition, whereas the C-terminal subunit (S2) mediates membrane fusion and host cell entry. The S1 domain itself contains a distinct N-terminal domain (NTD) followed by a receptor-binding domain (RBD) that is responsible for host cell recognition through interactions with the human cell-surface protein angiotensin-converting enzyme 2 (ACE2).

Most of the natural neutralizing antibodies (nAbs) that arise in response to SARS-CoV-2 infection target the S1 subunit, and while the most potent of these bind to the RBD and compete with ACE2 (Brouwer et al., 2020; Garrett Rappazzo et al., 2021; Tortorici et al., 2020), other RBD-binding antibodies are also neutralizing (Pinto et al., 2020). Cloning and recombinant production of these natural nAbs has led to the development of several drugs that have proven successful for inhibiting virus replication in patients and have been approved for therapy (Gottlieb et al., 2021; Razonable et al., 2021). Structural studies have shown that the most potent neutralizing IgGs bind to epitopes that overlap with the ACE2 epitope and take advantage of their bivalent nature to achieve high affinity by binding simultaneously to two of the three RBDs within the S-protein trimer (Miersch et al., 2021; Yan et al., 2021).

However, despite the significant clinical success of bivalent nAbs for treatment of COVID-19, they suffer from two significant drawbacks. First, even nAbs that are highly potent *in vitro* must be administered at extremely high doses, presumably at least in part due to low penetrance of IgGs to sites within the lungs where SARS-CoV-2 infects and replicates (Hart et al., 2001). Second, approved IgG drugs that were derived from nAbs that arose in response to the original strain of the virus have proven to be ineffective against many variants of concern (VoCs) that have arisen since. Moreover, there is some evidence that single agent nAb therapy may promote evolution of resistant variants (Jensen et al., 2021; Weisblum et al., 2020). Notably, most VoCs that resist current therapeutic nAbs contain mutations within the RBD, which disrupt binding to nAbs but not to ACE2 (Greaney et al., 2021; McCallum et al., 2021; Planas et al., 2021; Zhou et al., 2021).

These limitations on potency and breadth of coverage appear to be inherent to single-agent nAb drugs, but several options for alleviating these problems have been explored. Cocktails of two nAbs, ideally with non-overlapping epitopes, have been shown to be more potent and resistant to virus escape, but the considerable complexity and expense of combining two nAbs makes this approach problematic (Baum et al., 2020; Hansen et al., 2020; Weinreich et al., 2021). Alternatively, small modular Ab variable domains or non-antibody scaffolds have been assembled as multimers that can simultaneously engage all three RBDs on an S-protein trimer and thus enhance potency beyond bivalent binding (Cao et al., 2020; Hunt et al., 2021; Kayabolen et al., 2021; Walser et al., 2021; Xu et al., 2021). In another approach, a bispecific, bivalent IgG – with each arm targeting a different epitope on the RBD - has been shown to neutralize VoCs that escape from conventional IgGs (de Gasparo et al., 2020).

We previously explored approaches that expand the valency of conventional bivalent IgGs without introducing non-Ab scaffolds that may compromise drug development due to issues of immunogenicity, developability, and/or manufacturing. We recently isolated a human nAb (15033) against SARS-CoV-2 from a synthetic phage-displayed library, and engineered a variant nAb (15033-7) with changes in the light chain that enhanced potency (Miersch et al., 2021). We then constructed tetravalent nAbs by fusing additional copies of the 15033-7 Fab to the N- or C-terminus of the 15033-7 IgG, and we showed that these tetravalent nAbs exhibited enhanced potency against an original SARS-CoV-2 isolate and an evolved VoC. Importantly, the tetravalent nAbs could be produced at high yields with the same methods used for bivalent IgGs, without compromising biophysical properties required for effective drug development.

We now report a novel tetravalent Ab format that expands on our earlier work and provides the significant advantage of enabling facile, modular assembly of tetravalent, bi-paratopic nAbs containing two distinct paratopes targeting the SARS-CoV-2 RBD. To achieve this, we first converted the 15033-7 IgG (**Figure 1A**) into a diabody-Fc format (D-Fc, **Figure 1B**) consisting of VH and VL domains linked together in a single polypeptide, and we showed that the new format was comparable to the IgG format in terms of production, affinity, and potency. The homodimeric D-Fc format enabled the facile fusion of Fab arms to the C-terminus to generate tetravalent diabody-Fc-Fab (D-Fc-F) molecules (**Figure 1C**). Because the diabody paratope does not require a separate light chain, this format proved to be highly modular, and Fabs could be added without any issues of the Fab light chain interfering with the diabody or *vice versa*. Thus, we could easily add either the 15033-7 Fab or the Fab from a different nAb 15036 to assemble either a mono-paratopic or a bi-paratopic tetravalent D-Fc-F molecule, respectively. The bi-paratopic D-Fc-F could be produced with yields and purities similar to those of conventional IgGs and exhibited excellent biophysical properties comparable to those of antibody drugs. It neutralized a panel of pseudoviruses representing diverse SARS-CoV-2 VoCs with potencies that greatly exceeded those of the component IgGs and proved to be highly potent even against variants that resisted the IgGs and the mono-paratopic D-Fc-F. Most importantly, neutralization of both chimeric and authentic live VoCs in two independent laboratories validated these findings, confirming both high potency and breadth of coverage.

**Figure 1.**
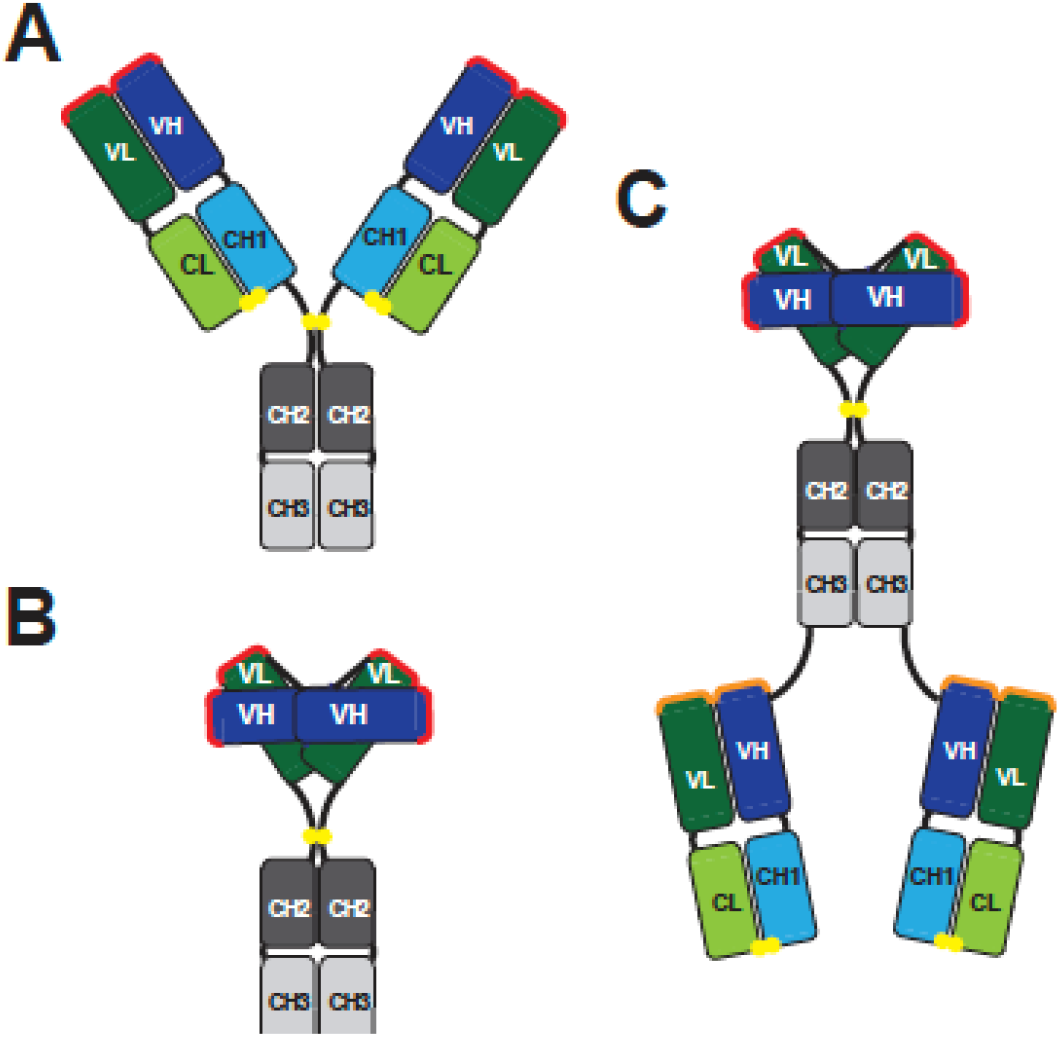
Antibody formats. Schematics are shown for **(A)** bivalent IgG, **(B)** bivalent diabody-Fc (D-Fc), and **(C)** bi-paratopic tetravalent diabody-Fc-Fab (D-Fc-F) formats. The paratopes in the IgG and D-Fc molecules are shown in red. In the D-Fc-F molecule, the diabody paratopes are shown in red and the Fab paratopes are shown in orange. Linkers are shown in black and intermolecular disulfide bonds are shown as yellow spheres. The domains are labeled and colored as follows: Fc, grey (CH2, dark; CH3, light); Fab heavy chain, blue (VH, dark; CH1, light); light chain, green (VL, dark; CL, light).

## Results

### Characterization of Ab 15036

To complement our previously characterized nAb 15033-7, we used well-established phage display methods with a highly validated synthetic human Fab-phage library (Persson et al., 2013) to identify a second Ab that bound to the RBD (see Materials and Methods). Ab 15036 had a very different paratope in comparison with 15033-7 (**Figure 2A**). Both Abs competed with each other for binding to immobilized RBD, as assessed by phage ELISAs in which binding of Fab-phage was measured in the presence or absence of saturating concentration of IgG protein (**Figure 2B**). However, whereas IgG 15033-7 completely blocked phage-displayed Fab 15036, IgG 15036 only partially blocked phage-displayed Fab 15033-7, suggesting that the epitopes for the two Fabs likely overlap to some extent but are not identical. Bilayer interferometry (BLI) assays showed that IgG 15036 bound to the S-protein trimer with a high affinity (apparent K_D_ = 220 pM) that was comparable to the previously determined affinity of IgG 15033-7 (apparent K_D_ = 70 pM) (**Figure 2C and Table 1**) (Miersch et al., 2021). Taken together, these results showed that Ab 15036 binds with high affinity to an epitope on the RBD that may overlap at least partially with the epitope of Ab 15033-7, a potent nAb that competes with ACE2. Thus, Ab 15036 should also be a potent nAb for preventing SARS-CoV-2 infection, which we confirmed below.

**Figure 2.**
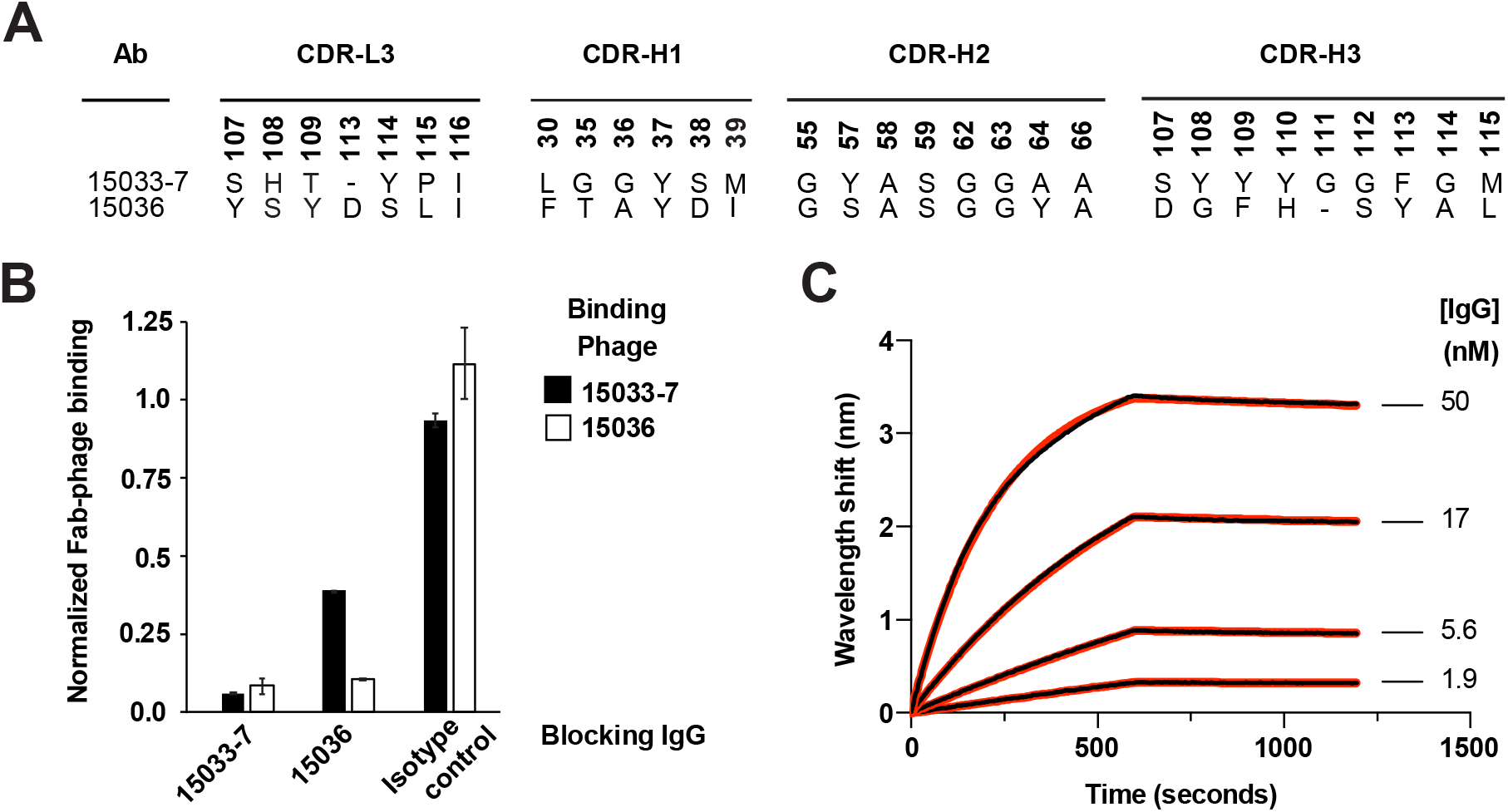
Characterization of antibody 15036. **(A)** CDR sequences of Abs 15033-7 and 15036. Positions are numbered according to the IMGT nomenclature (Lefranc et al., 2003). Only positions that were diversified in the Fab-phage library are shown. **(B)** Phage ELISAs for phage-displayed Fab 15033-7 (black bars) or Fab 15036 (white bars) binding to immobilized RBD in the presence of 250 nM blocking IgG 15033-7, 150336, or an isotype control. The binding signal was normalized to the signal in the absence of blocking IgG. **(C)** BLI sensor traces (black) for IgG 15036 binding to immobilized S-protein ECD trimer. Indicated concentrations of IgG 15036 were allowed to bind for 600 seconds and dissociation was monitored for an additional 600 seconds. The curves were globally fit (red) to a 1:1 binding model and derived binding constants are shown in **Table 1**.

**Table 1.**
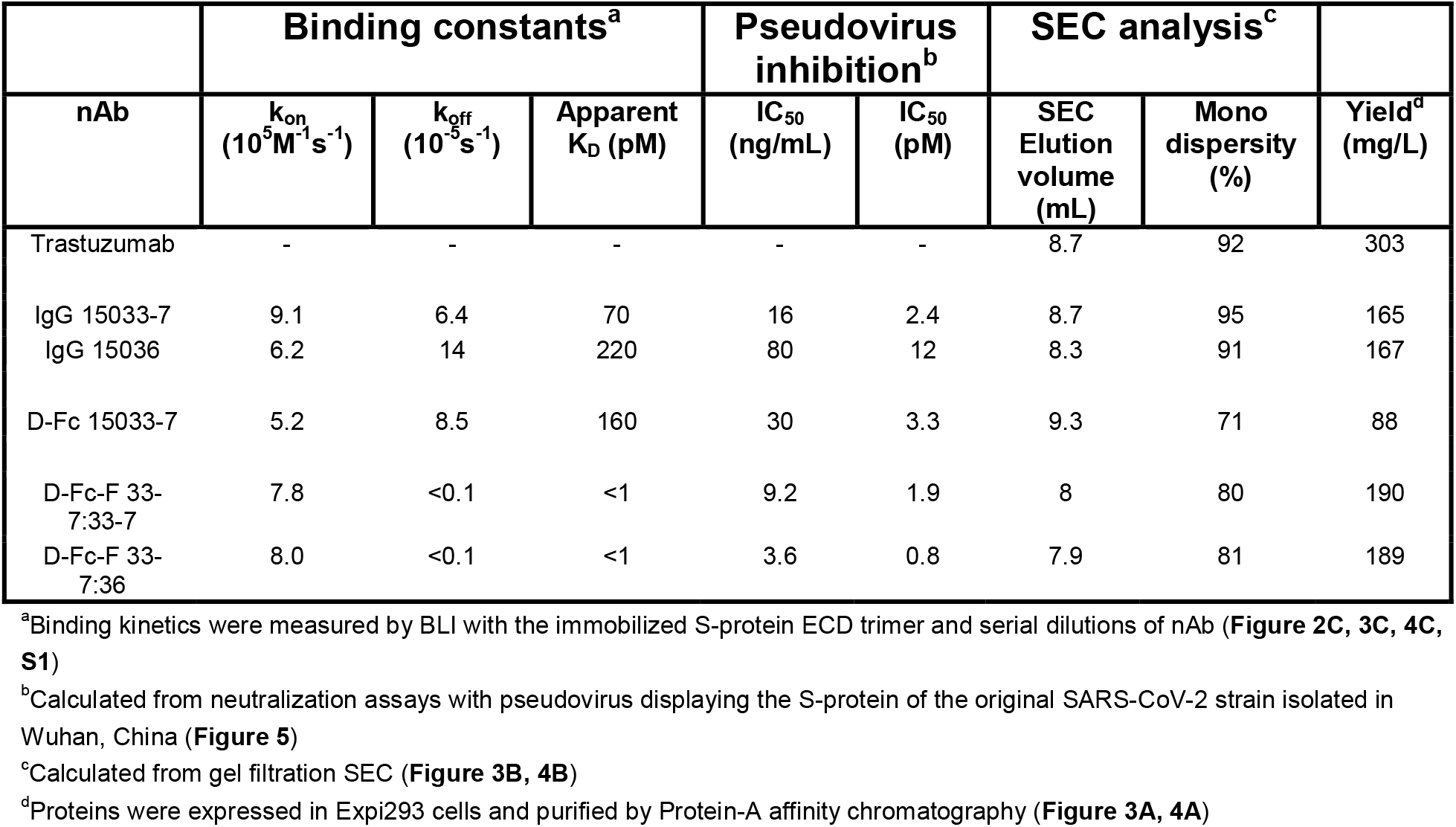
Summary of antibody characteristics.

| nAb | Binding constants<sup>a</sup> |  |  | Pseudovirus inhibition<sup>b</sup> |  | SEC analysis<sup>c</sup> |  | Yield<sup>d</sup><br>(mg/L) |
| --- | --- | --- | --- | --- | --- | --- | --- | --- |
| | $k_{on}$<br>($10^5 M^{-1} s^{-1}$) | $k_{off}$<br>($10^{-5} s^{-1}$) | Apparent<br>$K_D$ (pM) | IC<sub>50</sub><br>(ng/mL) | IC<sub>50</sub><br>(pM) | SEC<br>Elution<br>volume<br>(mL) | Mono<br>dispersity<br>(%) | |
| Trastuzumab | - | - | - | - | - | 8.7 | 92 | 303 |
| IgG 15033-7 | 9.1 | 6.4 | 70 | 16 | 2.4 | 8.7 | 95 | 165 |
| IgG 15036 | 6.2 | 14 | 220 | 80 | 12 | 8.3 | 91 | 167 |
| D-Fc 15033-7 | 5.2 | 8.5 | 160 | 30 | 3.3 | 9.3 | 71 | 88 |
| D-Fc-F 33-7:33-7 | 7.8 | <0.1 | <1 | 9.2 | 1.9 | 8 | 80 | 190 |
| D-Fc-F 33-7:36 | 8.0 | <0.1 | <1 | 3.6 | 0.8 | 7.9 | 81 | 189 |

<sup>a</sup>Binding kinetics were measured by BLI with the immobilized S-protein ECD trimer and serial dilutions of nAb (Figure 2C, 3C, 4C, S1)
<sup>b</sup>Calculated from neutralization assays with pseudovirus displaying the S-protein of the original SARS-CoV-2 strain isolated in Wuhan, China (Figure 5)
<sup>c</sup>Calculated from gel filtration SEC (Figure 3B, 4B)
<sup>d</sup>Proteins were expressed in Expi293 cells and purified by Protein-A affinity chromatography (Figure 3A, 4A)

### Characterization of D-Fc 15033-7

We produced IgG (**Figure 1A**) and D-Fc (**Figure 1B**) versions of Ab 15033-7 by transient transfection of Expi293F cells and purified the proteins by protein-A affinity chromatography. Both proteins were purified in high yield (165 or 88 mg/L for the IgG or D-Fc, respectively) (**Table 1**). Moreover, both proteins were highly pure, as assessed by SDS-PAGE (**Figure 3A**). The gels showed single bands of the appropriate size for intact molecules under non-reducing conditions with intact intermolecular disulfide bonds, whereas under reducing conditions without disulfide bonds, they showed a heavy-chain band and a light-chain band for the IgG, or a single band for a D-Fc monomer. Size exclusion chromatography (SEC) revealed that the IgG eluted as a predominantly (95%) monodisperse peak with an elution volume nearly identical to that of trastuzumab, whereas the D-Fc eluted as a predominantly (72%) monodisperse peak that was slightly delayed compared with trastuzumab, consistent with its smaller molecular weight (**Figure 3B**). Affinity measurements by BLI showed that the IgG and D-Fc proteins bound with similar high affinities to the trimeric S-protein ECD (apparent K_D_ = 70 or 160 pM, respectively) (**Figure 3C and Table 1**). Most importantly, in cell-based assays with a pseudovirus displaying the SARS-CoV-2 S-protein, the IgG and D-Fc proteins neutralized infection with essentially identical potencies (IC_50_ = 29 or 30 ng/mL, respectively) (**Figure 3D**).

**Figure 3.**
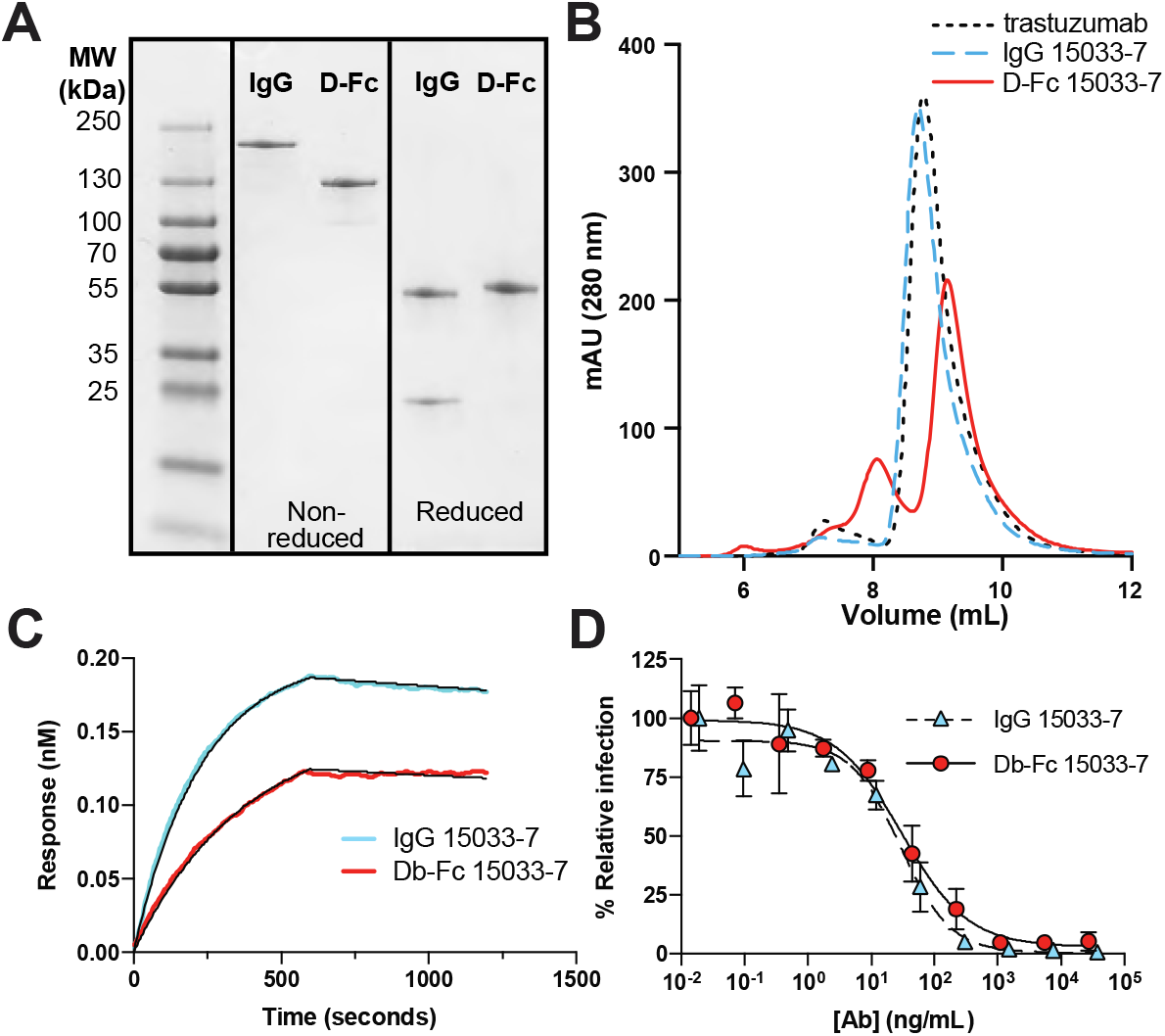
Characterization of D-Fc 15033-7. **(A)** SDS-PAGE _**300**_ analysis of IgG and D-Fc versions of Ab 15033-7 under non-reducing (left) or reducing conditions (right). **(B)** Analytical conditions (right). **(B)** Analytical gel filtration SEC of IgG 15033-7 (dashed blue), D-Fc 15033-7 conditions (right). **(B)** Analytical gel filtration SEC of IgG 15033-7 (dashed blue), D-Fc 15033-7 (solid red), and trastuzumab IgG (dotted black). **(C)** BLI sensor traces for 5 nM IgG 15033-7 (blue) or D-Fc 15033-7 (red) binding to immobilized S-protein ECD trimer. Abs were allowed to bind for 600 seconds, and dissociation was monitored for an additional 600 seconds. Traces and curve fits (black) for the complete analysis at various concentrations are shown in **Supplementary Figure S1** and the derived binding constants are shown in **Table 1. (D**) Neutralization of pseudovirus by IgG 15033-7 (blue triangles, dashed line) or D-Fc 15033-7 (red circles, solid line). Pseudovirus was generated with S-protein from the original strain isolated in Wuhan.

### Characterization of D-Fc-F proteins

We constructed D-Fc-F proteins (**Figure 1C**) in which the Fab of either the 15033-7 or 15036 nAb was fused to the C-terminus of D-Fc 15033-7. The resulting tetravalent molecules contained either four copies of the 15033-7 paratope (mono-paratopic, named D-Fc-F 33-7:33-7) or two copies of the 15033-7 paratope in the diabody head and two copies of the 15036 paratope in the Fab arms (bi-paratopic, named D-Fc-F 33-7:36). D-Fc-F 33-7:33-7 and 33-7:36 were both purified in similar high yields (190 or 189 mg/L, respectively) to high purity as evidenced by SDS-PAGE (**Figure 4A, Table 1**), with the same methods used for purification of IgG and D-Fc proteins. In gel filtration SEC, both proteins exhibited a predominant monodisperse peak (80% and 81% for D-Fc-F 33-7:33-7 or 33-7:36, respectively) that eluted earlier than the IgG trastuzumab, consistent with the larger size of the D-Fc-F protein (**Figure 4B and Table 1**). BLI analysis with trimeric S-protein ECD showed that D-Fc-F proteins bound very tightly and exhibited extremely slow off-rates that were beyond the dynamic range of the instrument, and consequently, apparent dissociation constants could not be measured accurately but were in the sub-picomolar range (apparent K_D_ < 1 pM) (**Figure 4C and Table 1**).

**Figure 4.**
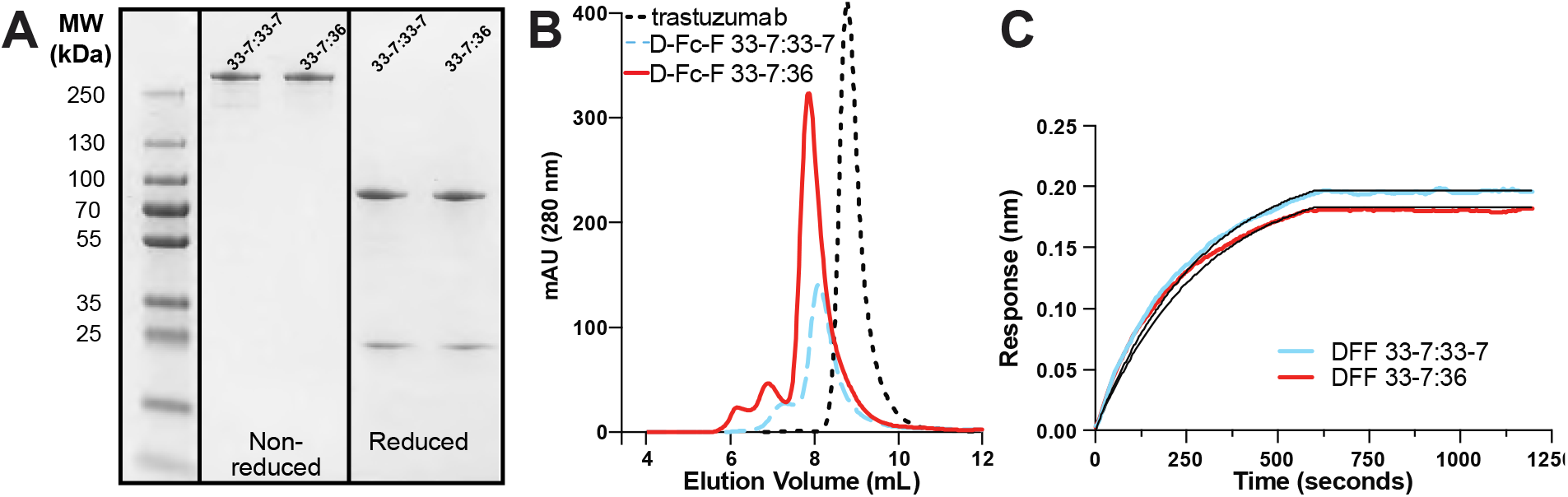
Characterization of D-Fc-F proteins. **(A)** SDS-PAGE analysis of D-Fc-F 33-7:33-7 and D-Fc-F 33-7:36 under non-reducing (left) or reducing conditions (right). **(B)** Analytical gel filtration SEC of D-Fc-F 33-7:33-7 (dashed blue), D-Fc-F 33-7:36 (solid red), and trastuzumab IgG (dotted black). **(C)** BLI sensor traces for 5 nM D-Fc-F 33-7:33-7 (blue) and D-Fc-F 33-7:36 (red) binding to immobilized S-protein ECD trimer. Abs were allowed to bind for 600 seconds, and dissociation was monitored for an additional 600 seconds. Traces and curve fits (black) for the complete analysis at various concentrations are shown in **Supplementary Figure S1** and the derived binding constants are shown in **Table 1**.

### Inhibition of virus infection in cell-based assays

To explore and compare the efficacy of the D-Fc-F proteins and their component IgG proteins for neutralization of SARS-CoV-2 VoCs, we first used mammalian cell infection assays with pseudoviruses consisting of HIV-gag-based, lentivirus-like particles pseudotyped with various SARS-CoV-2 S-proteins (Connor et al., 1995). We used a panel of seven pseudoviruses based on the S-protein of an early “wild-type” (WT) SARS-CoV-2 isolated in Washington (strain 2019 n-CoV/USA_WA1/2020) modified to incorporate RBD mutations corresponding to six VoCs, including strains that emerged in the United Kingdom (B.1.1.7), Italy (MB-61), South Africa (B.1.351), Brazil (P.1), India (B.1.617.1) and California (B.1.427/429) (See **Supplementary Table S1** for sequences). The two IgGs (15033-7 and 15036) and two D-Fc-Fs (33-7:33-7 and 33-7:33-6) were assayed against the pseudovirus panel (**Figure 5**) and IC_50_ values were calculated as a measure of potency (**Table 2**)

**Figure 5.**
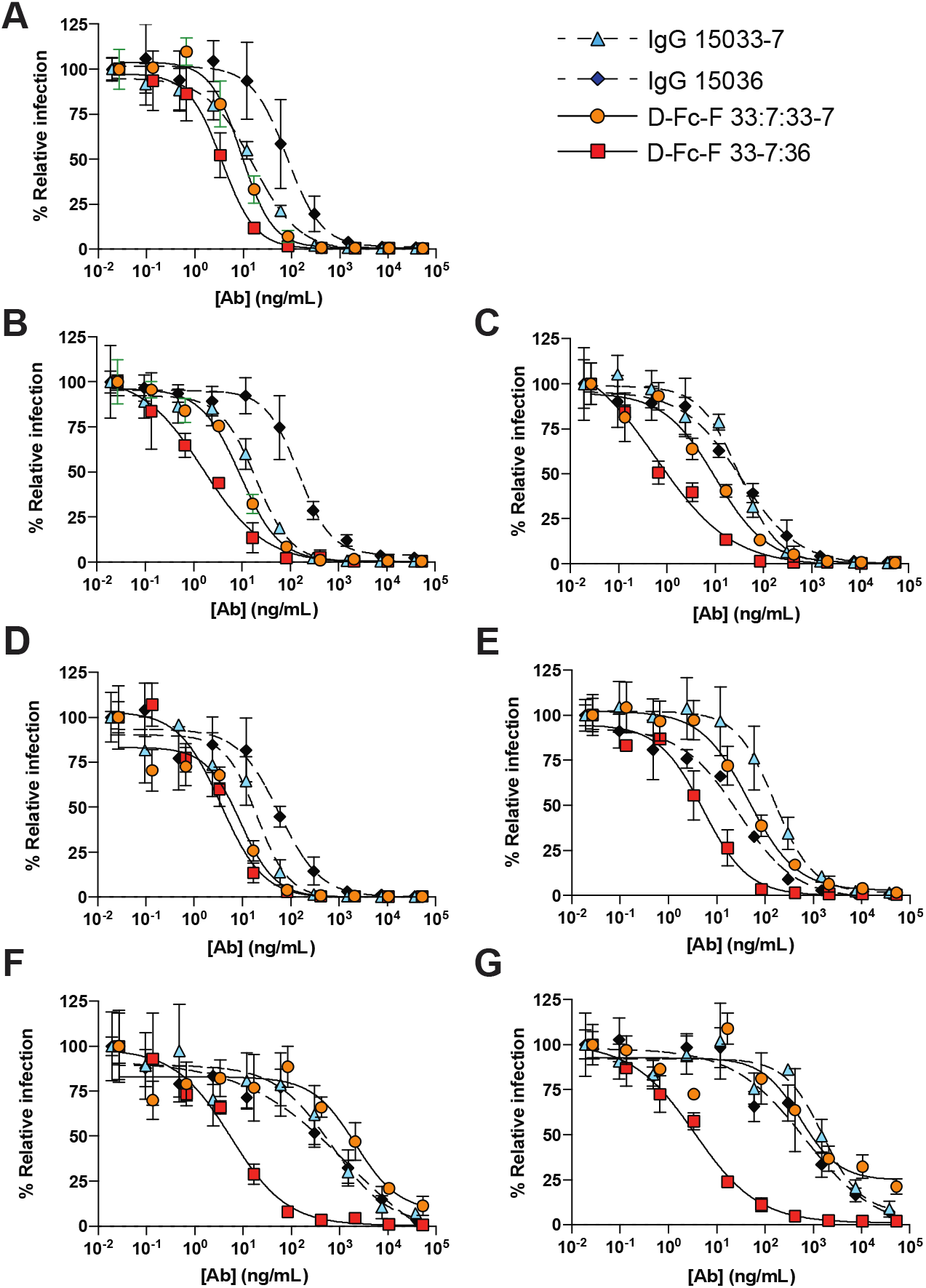
Neutralization of SARS-CoV-2 pseudoviruses. Neutralization assays were conducted with HIV-gag-based, lentivirus-like particles pseudotyped with the following SARS-CoV-2 S-proteins: **(A)** original strain from Wuhan, (**B**) Alpha (B.1.1.7) from the United Kingdom, (**C**) Kappa (B.1.617.1) from India, (**D**) Epsilon (B.1.427/429) from California, (**E**) MB-61 from Italy, (**F**) Beta (B.1.351) from South Africa, and (**G**) Gamma (P.1) from Brazil. The following nAbs were tested: IgG 15033-7 (cyan triangles, dashed lines), IgG 15036 (black diamonds, dashed lines), D-Fc-F 33-7:33-7 (orange circles, solid lines), and D-Fc-F 33-7:36 (red squares, solid lines). The pseudovirus was pre-treated with serial dilutions of nAb and infection of HEK293T cells stably expressing human ACE2 was measured relative to untreated control. Samples were run in duplicate, and results are representative of two independent experiments. Error bars indicate standard error of the mean. IC_50_ values were calculated from fit curves and are shown in **Table 2**.

**Table 2.** Summary of antibody IC_50_ values.

| Pseudovirus<sup>a</sup> |  |  | nAb IC<sub>50</sub> (ng/mL)<sup>b</sup> |  |  |  |
| --- | --- | --- | --- | --- | --- | --- |
| Origin | PANGO ID | Greek ID | IgG 15033-7 | IgG 15036 | D-Fc-F<br>33-7:33-7 | D-Fc-F<br>33-7:36 |
| Wuhan | - | - | 16 | 80 | 9.2 | 3.6 |
| Italy | MB-61 | - | 160 | 28 | 46 | 5.3 |
| UK | B.1.1.7 | Alpha | 20 | 150 | 9.2 | 1.6 |
| SA | B.1.351 | Beta | 680 | 1400 | 2100 | 5.6 |
| Manaus | P.1 | Gamma | 1600 | 680 | 550 | 3.5 |
| California | P.1.427/429 | Epsilon | 20 | 57 | 9.1 | 3.6 |
| India | B.1.617.1 | Kappa | 32 | 34 | 10 | 0.6 |

<sup>a</sup>Pseudovirus variants were generated by introducing RBD mutations corresponding to known variants of interest and variants of concern (Figure S2) to the plasmid encoding S-protein (original Wuhan strain) and used for pseudotyping.
<sup>b</sup>Values obtained by luminescent pseudovirus infection assay following exposure of ACE2-HEK293 cells to pseudovirus pre-incubated with serial dilutions of antibody.

IgG 15033-7 was highly effective against WT (**Figure 5A**, IC_50_ = 16 ng/mL), B.1.1.7 (**Figure 5B**, IC_50_ = 20 ng/mL), B.1.617.1 (**Figure 5C**, IC_50_ = 32 ng/mL) and B.1.427/429 (**Figure 5D**, IC_50_ = 20 ng/mL); moderately effective against MB-61 (**Figure 5E**, IC_50_ = 160 ng/mL); and ineffective against B.1.351 (**Figure 5F**, IC_50_ = 680 ng/mL) and P.1 (**Figure 5G**, IC_50_ = 1600 ng/mL). IgG 15036 exhibited a similar pattern of activities, but in general, was less potent than IgG 15033-7. The mono-paratopic, tetravalent D-Fc-F 33-7:33-7 was more potent against the five variants that were neutralized by the IgGs - WT, B.1.1.7, B.1.617.1, B.1.427/429 and MB-61 (**Figure 5A-E**, IC50 = 9-46 ng/mL) - but it also was ineffective against the two variants that resisted the IgGs - B.1.351 and P.1 (**Figure 5F-G**, IC_50_ = 2100 or 550 ng/mL, respectively). In contrast, the bi-paratopic, tetravalent D-Fc-F 33-7:36 was extremely potent against the entire pseudovirus panel (IC_50_ < 10 ng/mL) and was the most potent agent in every case (**Table 2**). Most dramatically, D-Fc-F 33-7:36 was extremely potent even against the variants that resisted the IgGs and D-Fc-F 33-7:33-7, and these assays showed >100-fold enhancements in some cases. Indeed, the IC_50_ values for D-Fc-F 33-7:36 against five of the seven pseudoviruses were virtually identical, suggesting that we may have reached the limits of the dynamic range for this assay and some potencies were underestimated.

Finally, and most importantly, we assessed the neutralization potency of D-Fc-F 33-7:36 in cell-based assays with authentic infectious virus in two independent laboratories. At Washington University in the USA, we used a panel of five chimeric viruses with S-protein variants representing diverse VoCs and observed extremely potent neutralization in all cases (IC_50_ = 1.5-7.2 ng/mL, **Figure 6A**). At the University of Brescia in Italy, we assessed activity against the currently dominant Delta variant and the highly recalcitrant Gamma variant with authentic virus, and we again observed extremely high potencies (IC_50_ = 8.5 or 12 ng/mL, respectively, **Figure 6B**). Taken together, these results confirmed that the bi-paratopic tetravalent D-Fc-F 33-7:36 is an extremely potent nAb for inhibition of SARS-CoV-2 infection with broad coverage against VoCs, including those that are resistant to approved nAb therapeutics.

**Figure 6.**
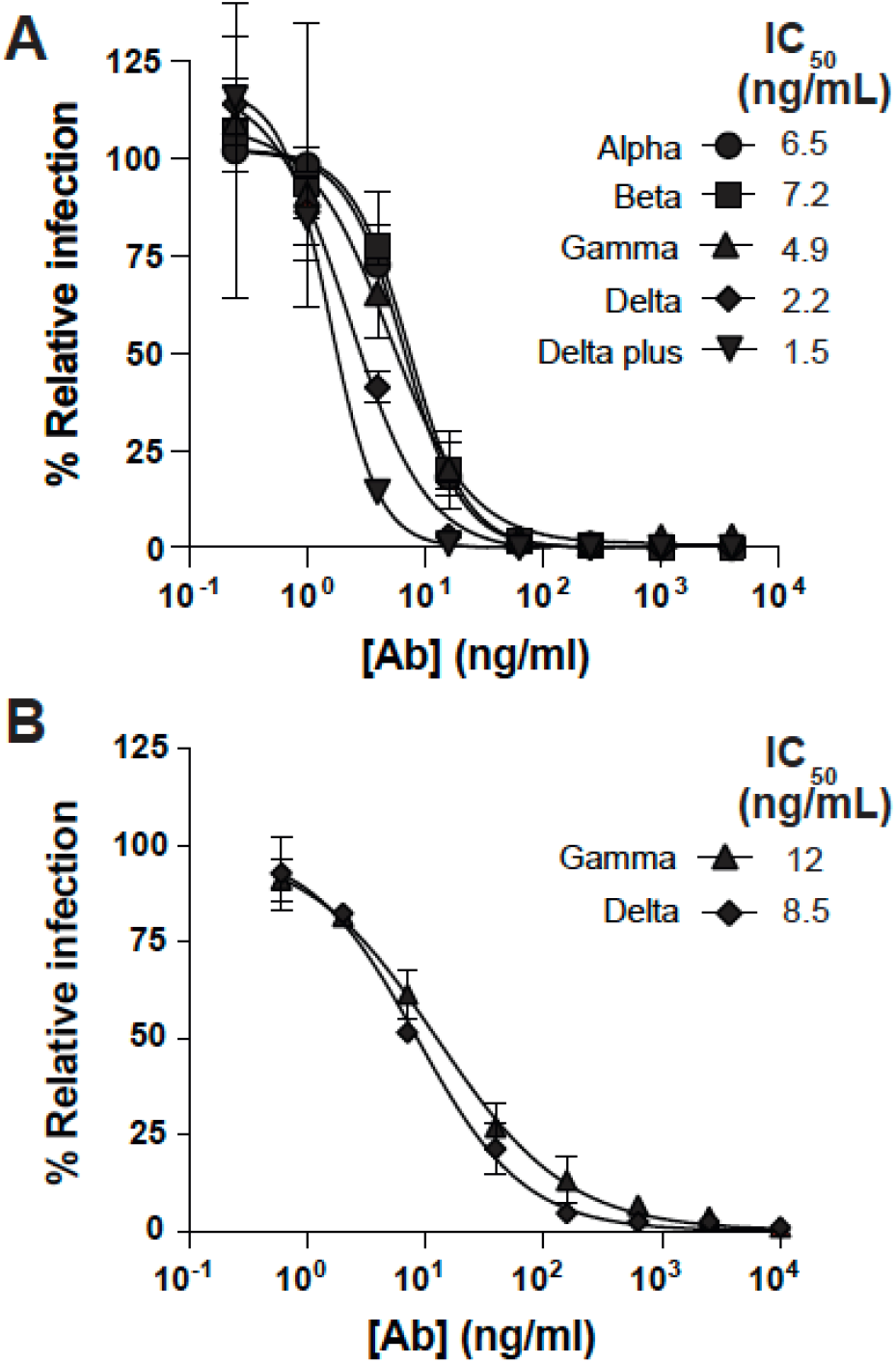
Neutralization of SARS-CoV-2 variants by D-Fc-F 33-7:36. **(A)** Neutralization of chimeric virus variants. A focus reduction test was used to evaluate infection of Vero E6 cells following exposure to chimeric virus variants pre-incubated with serial dilutions of D-Fc-F 33-7:36. Results shown are representative of two independent experiments. **(B)** Neutralization of authentic isolates of virus variants. A microneutralization assay was used to evaluate infection of Vero E6 cells following exposure to virus variants pre-incubated with serial dilutions of D-Fc-F 33-7:36. In both cases, samples were run in duplicate and normalized to infection in the absence of antibody with error bars shown as the standard deviation of the mean % relative infection values. Plots of nAb concentration versus relative infection for each were fit to estimate IC_50_ values shown in the legends.

## Discussion

Emergence of VoCs that resist approved nAbs (Hoffmann et al., 2021; Wang et al., 2021), as well as convalescent plasma and vaccine sera (Wall et al., 2021; Zhou et al., 2021), highlight the need for alternative therapeutics to meet this challenge. Modular and highly facile frameworks can fulfill this role as potential frontline therapies capable of neutralizing emerging VoCs and those that will emerge in the future. In particular, the E484K mutation found in the RBDs of Beta and Gamma VoCs – and also in the RBD of the Omicron variant (Bernasconi et al.) - impairs neutralization by bamlanivimab (Wang et al., 2021) and casirivimab (Kim et al., 2021; Wang et al., 2021), and the currently dominant Delta VoC also resists these nAbs to varying degrees (Planas et al., 2021).

The trimeric structure of the SARS-CoV-2 S-protein represents a significant vulnerability that can be exploited to enhance the potency of nAbs with advanced antibody engineering technologies. Numerous structural studies have shown that highly potent neutralizing IgGs take advantage of their two arms to bind simultaneously to two identical epitopes on the RBD, and bivalent binding enhances affinity and consequent potency of neutralization (Einav et al., 2019; Yan et al., 2021). We recently showed that fusing additional Fabs to a neutralizing IgG enhanced potency further and we also observed a concomitant improvement in neutralization of a VOC that resisted the IgG (Miersch et al., 2021). Moreover, structural studies of a tetravalent nAb in complex with the S-protein ECD confirmed that the two arms of the IgG bind simultaneously to two epitopes on the RBD, and intriguingly, suggested that a third arm can engage additional epitopes, within the same trimer or potentially a different trimer. These findings provide a molecular mechanism to explain the enhanced potency and breadth of coverage.

Here, we report a major advance in antibody engineering with a novel format that provides the crucial advantage of enabling rapid and facile combination of two distinct paratopes in a tetravalent nAb that retains the favorable biophysical properties of conventional IgG drugs. We started the engineering process by converting our potent neutralizing IgG 15033-7 (**Figure 1A**) to a D-Fc format (**Figure 1B**), which enabled modular attachment of additional Fabs at the C-terminus to produce tetravalent D-Fc-F proteins (**Figure 1C**). Importantly, the modular design strategy allowed for facile production of either the mono-paratopic D-Fc-F 33-7:33-7 or the bi-paratopic D-Fc-F 33-7:36, and both proteins proved to be comparable to IgGs in terms of biophysical properties (**Figure 4**). However, the tetravalent nAbs proved to be more potent than either IgG in neutralization assays with a panel of pseudoviruses representing SARC-CoV-2 VoCs (**Figure 5**). Most importantly, the bi-paratopic D-Fc-F 33-7:36 exhibited potent and broad neutralization of live viruses representing diverse VoCs, confirming that this tetravalent format represents an extremely promising means for the development of next-generation nAbs for the treatment of COVID-19.

Critically, our results revealed the value of bi-paratopic design, as D-Fc-F 33-7:36 was extremely potent against every VoC tested to date, including those that resisted both IgGs and even the mono-paratopic D-Fc-F 33-7:33-7. These results were not necessarily expected, given that the epitopes of nAbs 15033-7 and 15036 presumably overlap and exhibit similar activity patterns in the pseudovirus neutralization assays (**Figure 5**). Nonetheless, the two paratopes function differently in the tetravalent D-Fc-F context, as evidenced by the dramatic potency and breadth of coverage exhibited by D-Fc-F 33-7:36 compared with D-Fc-F 33-7:33-7. While the structural basis for the synergy between the two paratopes in D-Fc-F 33-7:36 remains to be determined, these results unequivocally establish the bi-paratopic, tetravalent D-Fc-F format as a powerful platform for next generation drug design to tackle SARS-CoV-2 and perhaps other viruses.

Indeed, our approach is truly agnostic and generally applicable to SARS-CoV-2 and to other viruses with similar construction. Importantly, *a priori* knowledge of structural information is not required to enable rapid testing of numerous paratope pairs in a high-throughput manner. We and other groups have already identified hundreds of nAbs targeting diverse neutralizing epitopes on the SARS-CoV-2 RBD and other regions of the S-protein (Cao et al., 2020b; Hansen et al., 2020; Liu et al., 2020; Miersch et al., 2021; Rogers et al., 2020; Shi et al., 2020; Wan et al., 2020; Wec et al., 2020), including broadly neutralizing nAbs (Garrett Rappazzo et al., 2021). Our D-Fc-F format allows modular combination of virtually any Fab with D-Fc 15033-7, thus enabling the facile production of hundreds of tetravalent nAbs that can tested systematically to develop the best possible frontline treatment for COVID-19 that can treat emerging VoCs and those that will emerge in the future.

## Acknowledgments

We thank Dr. James Rini for generously providing biotinylated S-protein and RBD.

## Author Contributions

^**†**^**Shane Miersch** Conceptualization, Methodology, Investigation, Supervision, Writing - Original Draft, Writing - Review & Editing, Visualization, Validation, Project administration

**Reza Saberianfar** Methodology, Investigation

**Chao Chen** Investigation

**Nitin Sharma** Investigation

**Gaya K. Amarasinghe** Investigation

**Arnaldo Caruso** Investigation

**Francesca Caccuri** Investigation

**Alberto Zani** Investigation

**Giuseppe Novelli** Investigation

^**\***^**§Sachdev Sidhu** Conceptualization, Writing - Original Draft, Supervision, Funding acquisition

## Materials

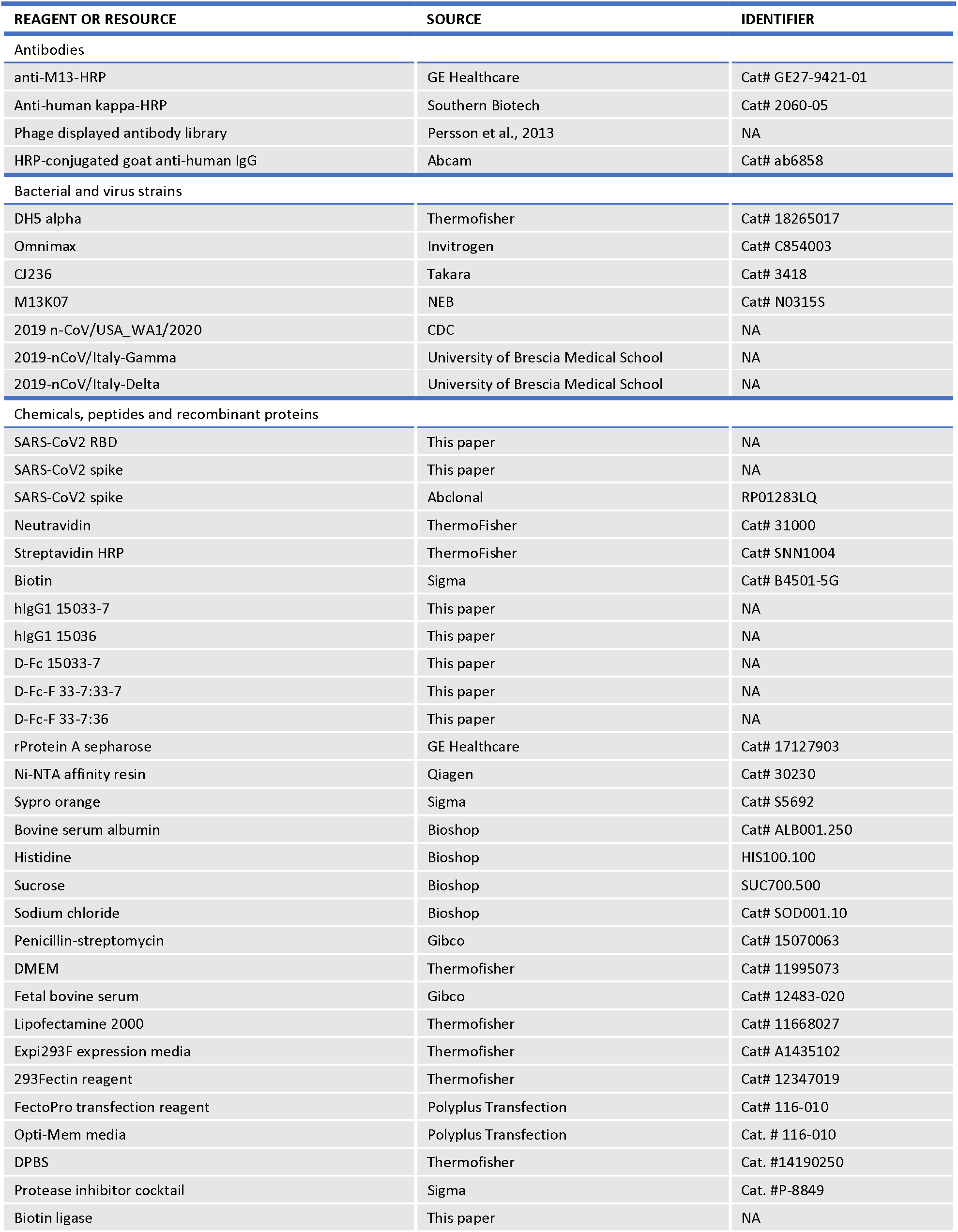
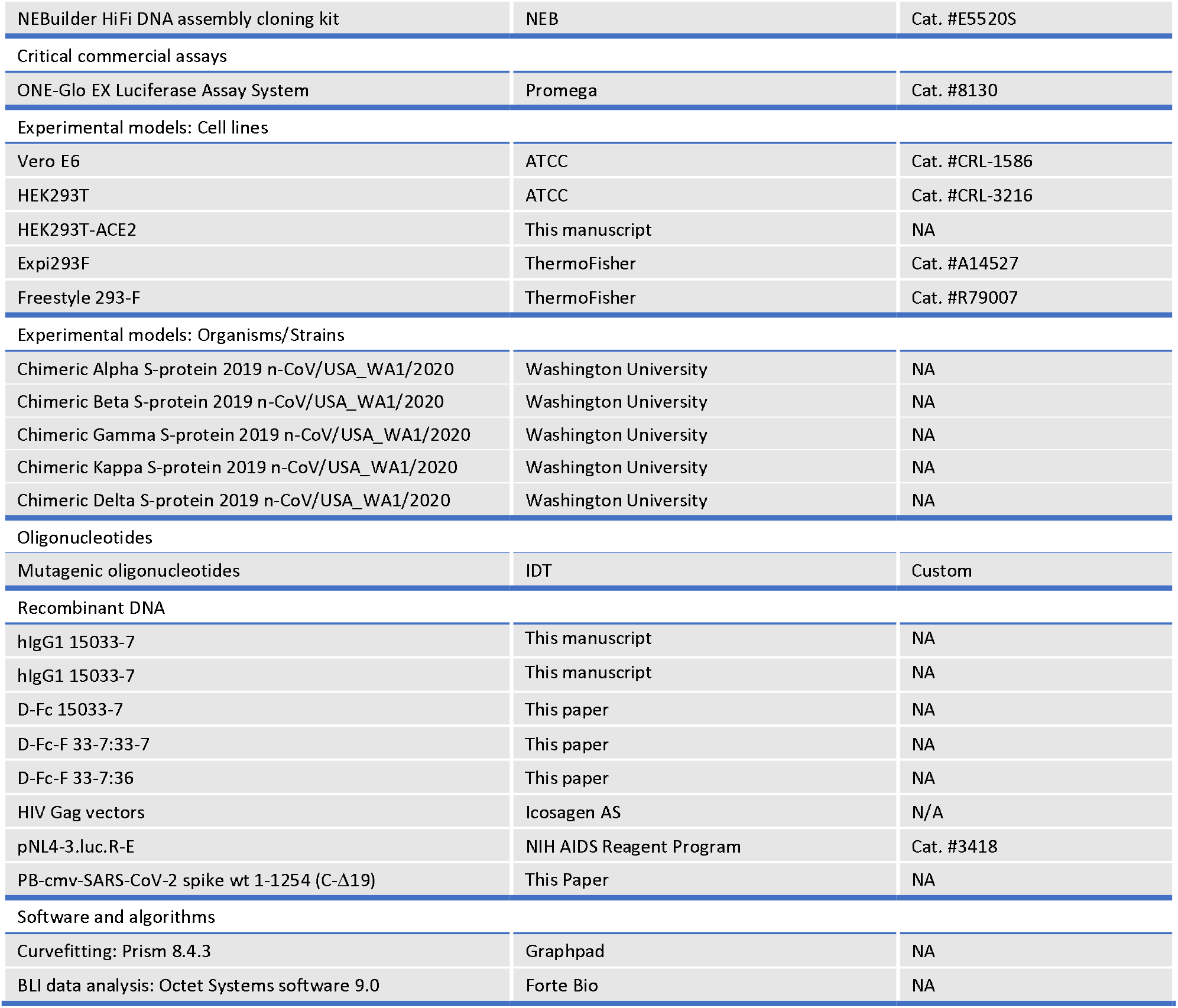

| REAGENT OR RESOURCE | SOURCE | IDENTIFIER |
| --- | --- | --- |
| Antibodies |  |  |
| anti-M13-HRP | GE Healthcare | Cat# GE27-9421-01 |
| Anti-human kappa-HRP | Southern Biotech | Cat# 2060-05 |
| Phage displayed antibody library | Persson et al., 2013 | NA |
| HRP-conjugated goat anti-human IgG | Abcam | Cat# ab6858 |
| Bacterial and virus strains |  |  |
| DH5 alpha | ThermoFisher | Cat# 18265017 |
| Omnimax | Invitrogen | Cat# C854003 |
| CJ236 | Takara | Cat# 3418 |
| M13K07 | NEB | Cat# N0315S |
| 2019 n-CoV/USA_WA1/2020 | CDC | NA |
| 2019-nCoV/Italy-Gamma | University of Brescia Medical School | NA |
| 2019-nCoV/Italy-Delta | University of Brescia Medical School | NA |
| Chemicals, peptides and recombinant proteins |  |  |
| SARS-CoV2 RBD | This paper | NA |
| SARS-CoV2 spike | This paper | NA |
| SARS-CoV2 spike | Abclonal | RP01283LQ |
| Neutravidin | ThermoFisher | Cat# 31000 |
| Streptavidin HRP | ThermoFisher | Cat# SNN1004 |
| Biotin | Sigma | Cat# B4501-5G |
| hlgG1 15033-7 | This paper | NA |
| hlgG1 15036 | This paper | NA |
| D-Fc 15033-7 | This paper | NA |
| D-Fc-F 33-7:33-7 | This paper | NA |
| D-Fc-F 33-7:36 | This paper | NA |
| rProtein A sepharose | GE Healthcare | Cat# 17127903 |
| Ni-NTA affinity resin | Qiagen | Cat# 30230 |
| Sypro orange | Sigma | Cat# S5692 |
| Bovine serum albumin | Bioshop | Cat# ALB001.250 |
| Histidine | Bioshop | HIS100.100 |
| Sucrose | Bioshop | SUC700.500 |
| Sodium chloride | Bioshop | Cat# SOD001.10 |
| Penicillin-streptomycin | Gibco | Cat# 15070063 |
| DMEM | ThermoFisher | Cat# 11995073 |
| Fetal bovine serum | Gibco | Cat# 12483-020 |
| Lipofectamine 2000 | ThermoFisher | Cat# 11668027 |
| Expi293F expression media | ThermoFisher | Cat# A1435102 |
| 293Fectin reagent | ThermoFisher | Cat# 12347019 |
| FectoPro transfection reagent | Polyplus Transfection | Cat# 116-010 |
| Opti-Mem media | Polyplus Transfection | Cat. # 116-010 |
| DPBS | ThermoFisher | Cat. #14190250 |
| Protease inhibitor cocktail | Sigma | Cat. #P-8849 |
| Biotin ligase | This paper | NA |
| NEBuilder HiFi DNA assembly cloning kit | NEB | Cat. #E5520S |
| Critical commercial assays |  |  |
| ONE-Glo EX Luciferase Assay System | Promega | Cat. #8130 |
| Experimental models: Cell lines |  |  |
| Vero E6 | ATCC | Cat. #CRL-1586 |
| HEK293T | ATCC | Cat. #CRL-3216 |
| HEK293T-ACE2 | This manuscript | NA |
| Expi293F | ThermoFisher | Cat. #A14527 |
| Freestyle 293-F | ThermoFisher | Cat. #R79007 |
| Experimental models: Organisms/Strains |  |  |
| Chimeric Alpha S-protein 2019 n-CoV/USA_WA1/2020 | Washington University | NA |
| Chimeric Beta S-protein 2019 n-CoV/USA_WA1/2020 | Washington University | NA |
| Chimeric Gamma S-protein 2019 n-CoV/USA_WA1/2020 | Washington University | NA |
| Chimeric Kappa S-protein 2019 n-CoV/USA_WA1/2020 | Washington University | NA |
| Chimeric Delta S-protein 2019 n-CoV/USA_WA1/2020 | Washington University | NA |
| Oligonucleotides |  |  |
| Mutagenic oligonucleotides | IDT | Custom |
| Recombinant DNA |  |  |
| hlgG1 15033-7 | This manuscript | NA |
| hlgG1 15033-7 | This manuscript | NA |
| D-Fc 15033-7 | This paper | NA |
| D-Fc-F 33-7:33-7 | This paper | NA |
| D-Fc-F 33-7:36 | This paper | NA |
| HIV Gag vectors | Icosagen AS | N/A |
| pNL4-3.luc.R-E | NIH AIDS Reagent Program | Cat. #3418 |
| PB-cmv-SARS-CoV-2 spike wt 1-1254 (C-Δ19) | This Paper | NA |
| Software and algorithms |  |  |
| Curvefitting: Prism 8.4.3 | Graphpad | NA |
| BLI data analysis: Octet Systems software 9.0 | Forte Bio | NA |

## Methods

### Cells

Mammalian cells were maintained in humidified environments at 37 °C in 5% CO_2_ in the indicated media. Vero E6 (ATCC, CRL-1586), HEK293T (ATCC, ACS-4500) and HEK293T cells stably overexpressing ACE2 were maintained at 37 °C in 5% CO_2_ in DMEM containing 10% (vol/vol) FBS. Expi293F cells (ThermoFisher, A14528) were maintained at 37 °C in 8% CO_2_ in Expi293F expression media (ThermoFisher, A1435101).

### Protein production and purification

The SARS-CoV-2 S-protein ECD was obtained as a kind gift from Dr. James Rini, produced and purified as described (Miersch et al., 2021) or was obtained commercially (Abclonal, RPO1283LQ). Purified proteins were site-specifically biotinylated in a reaction with 200 µM biotin, 500 µM ATP, 500 µM MgCl_2_, 30 µg/mL BirA, 0.1% (v/v) protease inhibitor cocktail and not more than 100 µM of the protein-AviTag substrate. The reactions were incubated at 30 °C for 2 hours and biotinylated proteins were purified by size-exclusion chromatography.

### Ab production and purification

IgG, D-Fc and D-Fc-F proteins were produced in Expi293F cells (ThermoFisher) by transient transfection, by diluting heavy and light chain expression plasmid DNA in OptiMem serum-free media (Gibco, 31985088) before the addition of and incubation with FectoPro (Polyplus Transfection, 10100007) for 10 minutes. For IgG and D-Fc-F production, equivalent amounts of plasmids encoding heavy chain or light chain were transfected, whereas for D-Fc production, a single construct encoding the D-Fc protein was transfected. Following addition of the DNA complex to Expi293F cells and a 5-day expression period, Abs were purified using rProtein-A Sepharose (GE Healthcare, 17127904), then buffer exchanged and concentrated using Amicon Ultra-15 Centrifugal Filter devices (Millipore, UFC805096). IgG and D-Fc proteins were stored in PBS (Gibco), and D-Fc-F proteins were stored in 10 mM L-histidine, 0.9% sucrose, 140 mM NaCl, pH 6.0.

### Phage display selections

A synthetic, phage-displayed Fab library (Persson et al., 2013b) was selected for binding to biotinylated SARS-CoV-2 RBD, as described (Miersch et al., 2021). Individual clones from the same selected phage pool that previously yielded Ab 15033 were screened by phage ELISA to identify additional RBD-binding Fab-phage clones, including the progenitor of Ab 15036. The heavy chain of the 15036 progenitor was combined with a library of light chains with diversity in CDR-L3, and Fab 15036 was isolated following additional rounds of selection, as described (Miersch et al., 2021).

### Phage ELISAs

Phage ELISAs were performed, as described (Miersch et al., 2021). Briefly, 96-well Nunc Maxisorp (Sigma-Aldrich, M9410) plates were coated with neutravidin and blocked with PBS, 0.2% BSA. Biotinylated target protein was captured from solution by incubation in neutravidin-coated and BSA-blocked wells for 15 minutes with shaking at room temperature. Wells were incubated for 1 hour with a saturating concentration of IgG in PBS, or with PBS alone, and subsequently, Fab-phage were added and incubated for 30 minutes. Plates were washed, incubated with an anti-M13 antibody-HRP conjugate (GE Healthcare, 27-9421-01), and developed with TMB substrate (KPL, KP-50-76-03), as described (Miersch et al., 2017).

### Construction of genes encoding D-Fc and D-Fc-F proteins

DNA fragments encoding heavy (VH) and light chain (VL) variable domains, (terminating at Ser^128^ or Lys^127^, respectively, IMGT numbering throughout (Lefranc et al., 2003)) were amplified by PCR from the IgG expression vectors. Db-Fc constructs were constructed using Gibson assembly (New England Biolabs, Ipswich, MA) to fuse these fragments with an intervening linker (sequence: GGGGS) with VH-linker-VL orientation, and then cloning the resultant insert into the pSCSTa mammalian expression vector, which resulted in the diabody being fused to the N-terminus of the hinge residue Asp^6^ with an intervening linker (sequence: GGGGSGGGGS).. D-Fc-F constructs were assembled by fusing a Fab heavy chain to Fc residue Gly^129^ at the C-terminus of the D-Fc with an intervening linker (sequence: GGGGSGGGGSGGGTG).

### Size exclusion chromatography

Protein samples (50 µg) were injected onto a TSKgel BioAssist G3SWxl column (Tosoh) fitted with a guard column using an NGC chromatography system and a C96 autosampler (Biorad). The column was preequilibrated in a PBS mobile phase and protein retention was monitored by absorbance at 215 nm during a 1.5 CV isocratic elution in PBS.

### Biolayer interferometry

The binding kinetics (k_on_ and k_off_) and apparent affinity (K_D_) of Abs binding to the S-protein ECD were determined by BLI with an Octet HTX instrument (ForteBio) at 1000 rpm and 25 °C. Biotinylated S-protein ECD was captured on streptavidin biosensors from a 2 µg/mL solution to achieve a binding response of 0.4-0.6 nm and unoccupied sites were quenched with 100 µg/mL biotin. Abs were diluted with assay buffer (PBS, 1% BSA, 0.05% Tween 20), and 67 nM of an irrelevant biotinylated protein of similar size was used as negative control. Following equilibration with assay buffer, loaded biosensors were dipped for 600 seconds into wells containing 3-fold serial dilutions of each Ab starting at 67 nM, and subsequently, were transferred back into assay buffer for 600 seconds. Binding response data were corrected by subtraction of response from a reference and were fitted with a 1:1 binding model using ForteBio Octet Systems software 9.0.

### Production of pseudoviruses

HEK-293 cells (ATCC) were seeded in a 6-well plate at 3 × 10^5^ cells/well in DMEM (ThermoFisher, 11995-065) supplemented with 10% FBS and 1% penicillin-streptomycin (Gibco, 15140122) and grown overnight at 37 °C with 5% CO_2_. HEK293 cells were co-transfected with 1 µg pNL4-3.luc.R-E-plasmid (luciferase expressing HIV-1 with defective envelop protein) (NIH AIDS Reagent Program, ARP2128) and 0.06 µg CMV-promoter driven plasmid encoding the S-protein using Lipofectamine™ 2000 transfection reagent (ThermoFisher, 11668027). Pseudovirus particles were harvested by collecting supernatant 48 hours after transfection and were filter sterilized (0.44 µm, Millipore-Sigma, SLHA033SS).

### Pseudovirus infection assays

HEK293T cells stably over-expressing full-length human ACE2 protein were seeded in 96-well white polystyrene microplates (Corning, CLS3610) at 3 × 10^4^ cells/well in DMEM (10% FBS and 1% penicillin-streptomycin) and were grown overnight at 37 °C with 5% CO_2_. Pseudovirus particles were mixed with Ab, incubated at room temperature for 10 minutes, and added to the cells. The cells were incubated at 37 °C with 5% CO_2_, the medium was replaced with fresh DMEM (10% FBS and 1% penicillin-streptomycin) after 6 hours, and again every 24 hours up to 72 hours. To measure the luciferase signal (pseudovirus entry), DMEM was removed and DPBS (ThermoFisher) was added to cells before mixing with an equal volume of ONE-Glo™ EX Luciferase Assay System (Promega, E8130). Relative luciferase units were measured using a BioTek Synergy Neo plate reader (BioTek Instruments Inc.). The data were analyzed by GraphPad Prism Version 8.4.3 (GraphPad Software, LLC).

### Authentic virus infection assays

To evaluate neutralization of SARS-Cov-2 VoCs, chimeric S-protein variants were generated on the genetic background of the Washington strain (2019 n-CoV/USA_WA1/2020) as described (Chen et al., 2021). Neutralization of these infectious chimeric VoCs was evaluated using a focus reduction neutralization test and potency was estimated as described (Miersch et al., 2021).

For neutralization of authentic VoCs, clinical isolates were propagated, sequenced to verify identity, then used in a micro-neutralization assay as described (Caccuri et al., 2020). In brief, serial four-fold dilutions of antibody, starting from 10 µg/mL, were pre-incubated with 10^2^ TCID_50_ of the virus per 100 µl at 37 °C for 1 hour. The antibody-virus mixture was transferred to 96-well tissue culture plates containing sub-confluent Vero E6 cells in duplicate and incubated at 37 °C and 5% CO_2_ for 4 days before washing, fixation, staining and being read by a spectrophotometer at 590 nm as described (Manenti et al., 2020).

## REFERENCES

Baum, A., Fulton, B.O., Wloga, E., Copin, R., Pascal, K.E., Russo, V., Giordano, S., Lanza, K., Negron, N., Ni, M., et al. (2020). Antibody cocktail to SARS-CoV-2 spike protein prevents rapid mutational escape seen with individual antibodies. Science 369, 1014–1018.

Bernasconi, A., Pinoli, P., al Khalaf, R., Alfonsi, T., Canakoglu, A., Cilibrasi, L., and Ceri, S. Report on Omicron Spike mutations on epitopes and immunological/epidemiological/kinetics effects from literature - SARS-CoV-2 coronavirus / nCoV-2019 Genomic Epidemiology - Virological.

Brouwer, P.J.M., Caniels, T.G., van der Straten, K., Snitselaar, J.L., Aldon, Y., Bangaru, S., Torres, J.L., Okba, N.M.A., Claireaux, M., Kerster, G., et al. (2020). Potent neutralizing antibodies from COVID-19 patients define multiple targets of vulnerability. Science (New York, N.Y.) 369, 643–650.

Cao, L., Goreshnik, I., Coventry, B., Case, J.B., Miller, L., Kozodoy, L., Chen, R.E., Carter, L., Walls, A.C., Park, Y.J., et al. (2020). De novo design of picomolar SARS-CoV-2 miniprotein inhibitors. Science 370.

Connor, R.I., Chen, B.K., Choe, S., and Landau, N.R. (1995). Vpr Is Required for Efficient Replication of Human Immunodeficiency Virus Type-1 in Mononuclear Phagocytes. Virology 206, 935–944.

Einav, T., Yazdi, S., Coey, A., and Bjorkman, P.J. (2019). Harnessing Avidity: Quantifying the Entropic and Energetic Effects of Linker Length and Rigidity for Multivalent Binding of Antibodies to HIV-1. Cell Systems 9, 466-474.e7.

Garrett Rappazzo, C., Tse, L. v., Kaku, C.I., Wrapp, D., Sakharkar, M., Huang, D., Deveau, L.M., Yockachonis, T.J., Herbert, A.S., Battles, M.B., et al. (2021). Broad and potent activity against SARS-like viruses by an engineered human monoclonal antibody. Science (New York, N.Y.) 371, 823–829.

de Gasparo, R., Pedotti, M., Simonelli, L., Nickl, P., Muecksch, F., Cassaniti, I., Percivalle, E., C Lorenzi, J.C., Mazzola, F., Magrì, D., et al. (2020). Bispecific IgG neutralizes SARS-CoV-2 variants and prevents escape in mice. Nature 593.

Gottlieb, R.L., Nirula, A., Chen, P., Boscia, J., Heller, B., Morris, J., Huhn, G., Cardona, J., Mocherla, B., Stosor, V., et al. (2021). Effect of Bamlanivimab as Monotherapy or in Combination With Etesevimab on Viral Load in Patients With Mild to Moderate COVID-19: A Randomized Clinical Trial. JAMA 325, 632–644.

Greaney, A.J., Starr, T.N., Gilchuk, P., Zost, S.J., Binshtein, E., Loes, A.N., Hilton, S.K., Huddleston, J., Eguia, R., Crawford, K.H.D., et al. (2021). Complete Mapping of Mutations to the SARS-CoV-2 Spike Receptor-Binding Domain that Escape Antibody Recognition. Cell Host & Microbe 29, 44-57.e9.

Hansen, J., Baum, A., Pascal, K.E., Russo, V., Giordano, S., Wloga, E., Fulton, B.O., Yan, Y., Koon, K., Patel, K., et al. (2020). Studies in humanized mice and convalescent humans yield a SARS-CoV-2 antibody cocktail. Science (New York, N.Y.) 369, 1010–1014.

Hart, T.K., Sellers, T.S., Maleeff, B.E., Eustis, S., Schwartz, L.W., Bugelski, P.J., Herzyk, D.J., Cook, R.M., Zia-Amirhosseini, P., Minthorn, E., et al. (2001). Preclinical efficacy and safety of mepolizumab (SB-240563), a humanized monoclonal antibody to IL-5, in cynomolgus monkeys. Journal of Allergy and Clinical Immunology 108, 250–257.

Hoffmann, M., Kleine-Weber, H., Schroeder, S., Mü, M.A., Drosten, C., and Pö, S. (2020). SARS-CoV-2 Cell Entry Depends on ACE2 and TMPRSS2 and Is Blocked by a Clinically Proven Protease Inhibitor. Cell 181, 271-280.e8.

Hoffmann, M., Hofmann-Winkler, H., Krüger, N., Kempf, A., Nehlmeier, I., Graichen, L., Arora, P., Sidarovich, A., Moldenhauer, A.S., Winkler, M.S., et al. (2021). SARS-CoV-2 variant B.1.617 is resistant to bamlanivimab and evades antibodies induced by infection and vaccination. Cell Reports 36.

Hunt, A.C., Case, J.B., Park, Y.-J., Cao, L., Wu, K., Walls, A.C., Liu, Z., Bowen, J.E., Yeh, H.-W., Saini, S., et al. (2021). Multivalent designed proteins protect against SARS-CoV-2 variants of concern. BioRxiv.

Jensen, B., Luebke, N., Feldt, T., Keitel, V., Brandenburger, T., Kindgen-Milles, D., Lutterbeck, M., Freise, N.F., Schoeler, D., Haas, R., et al. (2021). Emergence of the E484K mutation in SARS-COV-2-infected immunocompromised patients treated with bamlanivimab in Germany. The Lancet Regional Health – Europe 8, 100164.

Kayabolen, A., Akcan, U., Ozturan, D., Sarayloo, E., Nurtop, E., Ozer, B., Sahin, G.N., Dogan, O., Lack, N., Kaya, M., et al. (2021). Protein scaffold-based multimerization of soluble ACE2 efficiently blocks SARS-CoV-2 infection in vitro. BioRxiv 2021.01.04.425128.

Kim, Y.J., Jang, U.S., Soh, S.M., Lee, J.Y., and Lee, H.R. (2021). The Impact on Infectivity and Neutralization Efficiency of SARS-CoV-2 Lineage B.1.351 Pseudovirus. Viruses 13.

Klein, S., Cortese, M., Winter, S.L., Wachsmuth-Melm, M., Neufeldt, C.J., Cerikan, B., Stanifer, M.L., Boulant, S., Bartenschlager, R., and Chlanda, P. (2020). SARS-CoV-2 structure and replication characterized by in situ cryo-electron tomography. BioRxiv 2020.06.23.167064.

McCallum, M., Bassi, J., Marco, A. de, Chen, A., Walls, A.C., Iulio, J. di, Tortorici, M.A., Navarro, M.-J., Silacci-Fregni, C., Saliba, C., et al. (2021). SARS-CoV-2 immune evasion by the B.1.427/B.1.429 variant of concern. Science 373, 648–654.

Miersch, S., Li, Z., Saberianfar, R., Ustav, M., Brett Case, J., Blazer, L., Chen, C., Ye, W., Pavlenco, A., Gorelik, M., et al. (2021). Tetravalent SARS-CoV-2 Neutralizing Antibodies Show Enhanced Potency and Resistance to Escape Mutations. Journal of Molecular Biology 433, 167177.

Persson, H., Ye, W., Wernimont, A., Adams, J.J., Koide, A., Koide, S., Lam, R., and Sidhu, S.S. (2013). CDR-H3 diversity is not required for antigen recognition by synthetic antibodies. Journal of Molecular Biology 425, 803–811.

Pinto, D., Park, Y.J., Beltramello, M., Walls, A.C., Tortorici, M.A., Bianchi, S., Jaconi, S., Culap, K., Zatta, F., de Marco, A., et al. (2020). Cross-neutralization of SARS-CoV-2 by a human monoclonal SARS-CoV antibody. Nature 2020 583:7815 583, 290–295.

Planas, D., Veyer, D., Baidaliuk, A., Staropoli, I., Guivel-Benhassine, F., Rajah, M.M., Planchais, C., Porrot, F., Robillard, N., Puech, J., et al. (2021). Reduced sensitivity of SARS-CoV-2 variant Delta to antibody neutralization. Nature 2021 596:7871 596, 276–280.

Razonable, R.R., Pawlowski, C., O’Horo, J.C., Arndt, L.L., Arndt, R., Bierle, D.M., Borgen, M.D., Hanson, S.N., Hedin, M.C., Lenehan, P., et al. (2021). Casirivimab–Imdevimab treatment is associated with reduced rates of hospitalization among high-risk patients with mild to moderate coronavirus disease-19. EClinicalMedicine 40, 101102.

Tortorici, M.A., Beltramello, M., Lempp, F.A., Pinto, D., Dang, H. v., Rosen, L.E., McCallum, M., Bowen, J., Minola, A., Jaconi, S., et al. (2020). Ultrapotent human antibodies protect against SARS-CoV-2 challenge via multiple mechanisms. Science (New York, N.Y.) 370, 950–957.

Wall, E.C., Wu, M., Harvey, R., Kelly, G., Warchal, S., Sawyer, C., Daniels, R., Hobson, P., Hatipoglu, E., Ngai, Y., et al. (2021). Neutralising antibody activity against SARS-CoV-2 VOCs B.1.617.2 and B.1.351 by BNT162b2 vaccination. Lancet (London, England) 397, 2331–2333.

Walls, A.C., Park, Y.J., Tortorici, M.A., Wall, A., McGuire, A.T., and Veesler, D. (2020). Structure, Function, and Antigenicity of the SARS-CoV-2 Spike Glycoprotein. Cell 181, 281-292.e6.

Walser, M., Rothenberger, S., Hurdiss, D.L., Schlegel, A., Calabro, V., Fontaine, S., Villemagne, D., Paladino, M., Hospodarsch, T., Neculcea, A., et al. (2021). Highly potent anti-SARS-CoV-2 multivalent DARPin therapeutic candidates. BioRxiv 2020.08.25.256339.

Wang, P., Nair, M.S., Liu, L., Iketani, S., Luo, Y., Guo, Y., Wang, M., Yu, J., Zhang, B., Kwong, P.D., et al. (2021). Antibody resistance of SARS-CoV-2 variants B.1.351 and B.1.1.7. Nature 2021 593:7857 593, 130–135.

Weinreich, D.M., Sivapalasingam, S., Norton, T., Ali, S., Gao, H., Bhore, R., Xiao, J., Hooper, A.T., Hamilton, J.D., Musser, B.J., et al. (2021). REGEN-COV Antibody Combination and Outcomes in Outpatients with Covid-19. New England Journal of Medicine.

Weisblum, Y., Schmidt, F., Zhang, F., DaSilva, J., Poston, D., Lorenzi, J.C.C., Muecksch, F., Rutkowska, M., Hoffmann, H.H., Michailidis, E., et al. (2020). Escape from neutralizing antibodies 1 by SARS-CoV-2 spike protein variants. ELife 9, 1.

Xu, J., Xu, K., Jung, S., Conte, A., Lieberman, J., Muecksch, F., Lorenzi, J.C.C., Park, S., Schmidt, F., Wang, Z., et al. (2021). Nanobodies from camelid mice and llamas neutralize SARS-CoV-2 variants. Nature 2021 595:7866 595, 278–282.

Yan, R., Wang, R., Ju, B., Yu, J., Zhang, Y., Liu, N., Wang, J., Zhang, Q., Chen, P., Zhou, B., et al. (2021). Structural basis for bivalent binding and inhibition of SARS-CoV-2 infection by human potent neutralizing antibodies. Cell Research 2021 31:5 31, 517–525.

Zhou, D., Dejnirattisai, W., Supasa, P., Liu, C., Mentzer, A.J., Ginn, H.M., Zhao, Y., Duyvesteyn, H.M.E., Tuekprakhon, A., Nutalai, R., et al. (2021). Evidence of escape of SARS-CoV-2 variant B.1.351 from natural and vaccine-induced sera. Cell 184, 2348-2361.e6.

