## Supplemental Information for "Ultrapotent and Broad Neutralization of SARS-CoV-2 Variants by Modular, Tetravalent, Bi-paratopic Antibodies"

### Supplementary Figures and Figure Legends

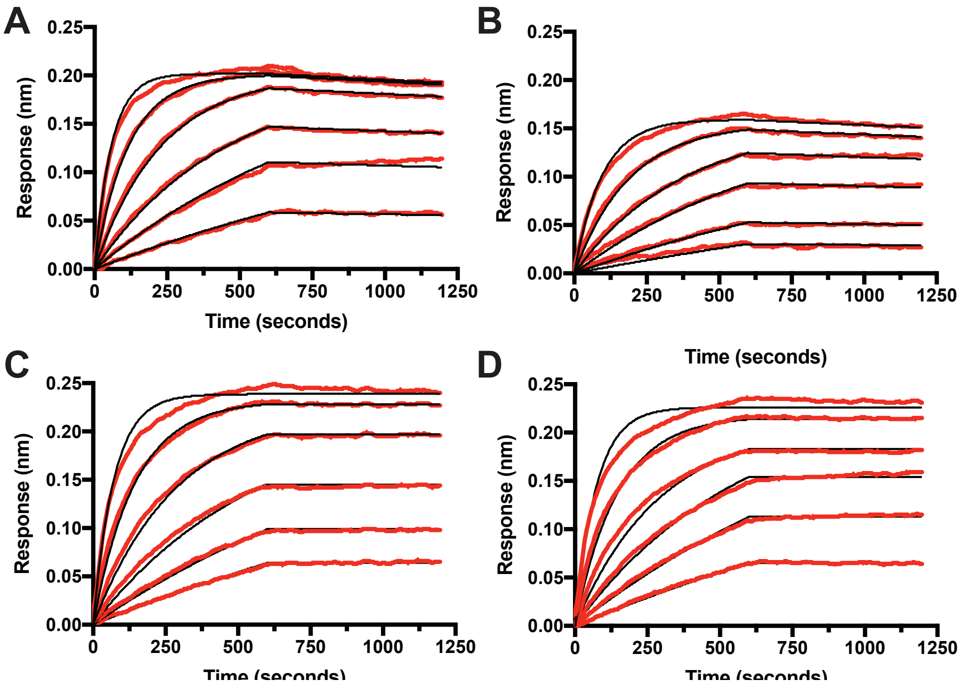


**Figure S1. Binding sensorgrams for nAb:S-protein interactions.** Binding signals (red) of nAb-S-protein interactions were determined for **A)** IgG 15033-7, **B)** D-Fc 15033-7, **C)** D-Fc-F 33-7:33-7 and **D)** D-Fc-F 33-7:36 by biolayer interferometry exposing serial 2-fold nAb dilutions (from 20 nM) to immobilized sensor-immobilized S-protein. Global fits (black) were obtained using a 1:1 binding model.

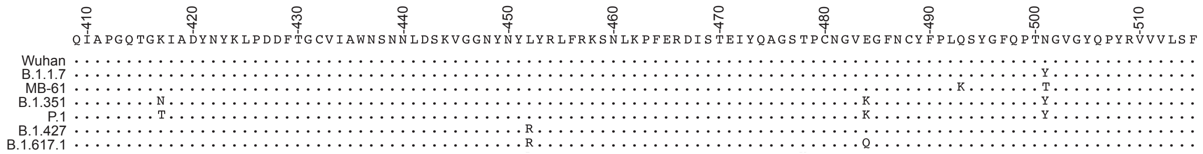


**Figure S2 – S-protein variant mutations used to generate pseudovirus variants** A segment of the RBD of the SARS-COV-2 S-protein corresponding to residues 409-515 showing the mutations introduced in to the RBD to generate pseudovirus for infection assays (**Fig. 5**).
